# Specific T cell factors exist

**DOI:** 10.1101/100024

**Authors:** Geoffrey W. Hoffmann, Reginald M. Gorczynski

## Abstract

The symmetrical immune network theory has been developed since 1975 and is based on the existence of specific T cell factors. The existence of specific T cell factors is controversial. We confirm the existence of these specific immune system components by demonstrating that a rigorous 1975 experiment by Takemori and Tada can be reproduced. We also briefly review how specific T cell factors play a role in the induction of specific tolerance and in the induction of immunity according to the symmetrical immune network theory.

## Introduction

The symmetrical immune network theory has been developed in peer-reviewed papers (Hoffmann et al., 1986, Hoffmann, 1994), and non-peer-reviewed publications (Hoffmann, 2008; Hoffmann, 2010). The theory is based on the presumed existence of specific T cell factors. Many immunologists documented the existence of specific T cell factors in the 1970s and 1980s (Nelson, 1970; Evans et al., 1972; Takemori and Tada, 1975, Sy et al., 1979, Hirai and Nisonoff, 1980; Shiozawa et al., 1980, 1984; Tada 1984). These were detected in various assays, including assays that showed specific help (Shiozawa et al, 1980, 1984) or specific suppression (Takemori and Tada, 1975, Hirai and Nisonoff, 1980, Tada 1984) of immune responses. The existence of these molecules is nevertheless controversial (Melchers, 1991, Varela and Coutinho, 1991), and they are not mentioned in current immunology textbooks, including for example the widely-used Kuby Immunology, Seventh Edition (Owen et al. 2013).

In 1991 one of us pointed out that the reproducibility of an experiment by Takemori and Tada (1975) is a suitable criterion for the existence of these factors (Hoffmann, 1991). We argued that if that experiment could not be reproduced the symmetrical immune network theory was an inadequate explanation of immunoregulation. On the other hand, if the experiment could be reproduced, this would imply that such factors do indeed exist, and their existence is currently being erroneously widely ignored. To our knowledge, in the 25 years since 1991, no-one has claimed to be unable to reproduce the Takemori-Tada result, nor has anyone published confirmation that the experiment is reproducible.

## Experiment

BALB/c mice were immunized twice with keyhole limpet hemocyanin (KLH) or bovine gamma globulin (BGG) on days 0 and 14. The mice were sacrificed at day 28 and thymus cells from the mice were sonicated. Extracts were used to attenuate anti-DNP IgG responses of mice of the same strain challenged with DNP-KLH or DNP-BGG. The experiment was performed with reciprocal specificity controls, and hence rigorously demonstrated specificity of the factors.

## Results

Data shown in Fig 1 are from an experiment designed to reproduce the Takemori and Tada 1975 finding of specific T cell factors. Mice were challenged with DNP-BGG or DNP-KLH following pre-treatment with lymphocyte extracts derived from animals immunized with carrier (KLH or BGG) alone. The results demonstrate highly specific suppression with reciprocal specificity in agreement with the Takemori and Tada 1975 result.

**Legend to Fig 1:**
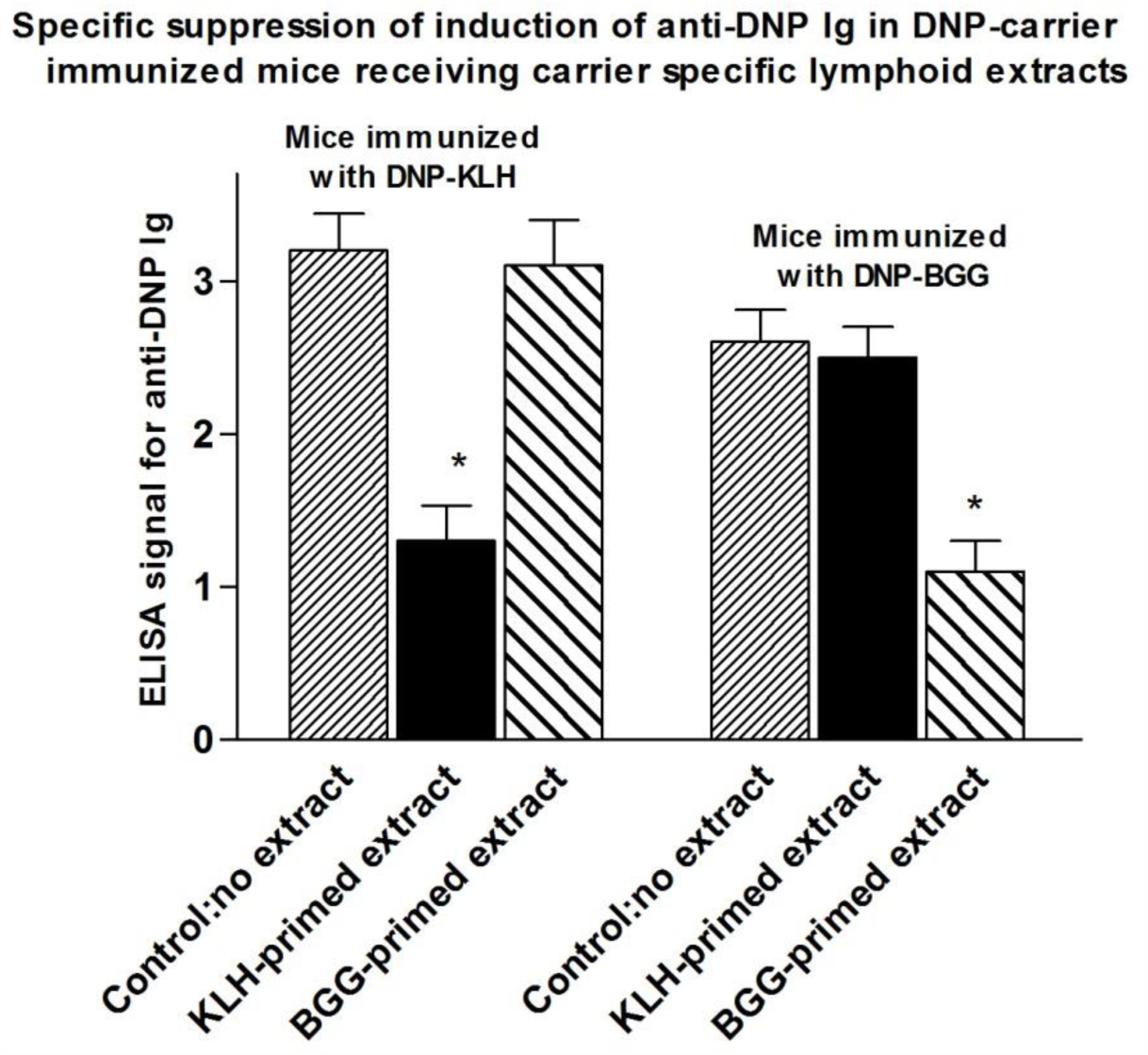
Cell free extract was obtained from sonicated (spleen + thymocyte) cell preparations from mice primed twice on days 0 and 14 with BGG or KLH. The extract was infused into mice that subsequently received DNP coupled to BGG or DNP coupled to KLH. IgG responses to DNP were measured. * indicates p<0.05 compared with no extract control (ANOVA).

## The postulated role of specific T cell factors in an immune response

The symmetrical immune network theory involves symmetrical stimulatory, inhibitory and killing interactions (Hoffmann, 2008). A simple symmetrical network model that includes antigen (Ag), antigen-specific lymphocytes (T+ and B+), antiidiotypic lymphocytes (T− and B−) and non-specific accessory (A) cells is shown in Figure 2. A cells include monocytes, macrophages and dendritic cells. In the model an immune response involves an immunogenic form of an antigen stimulating T+ and B+ cells, specific T cell factors from the T+ cells binding to a receptor on the surface of A cells, and the antigen activating the A cell by cross-linking a receptor on the A cell via the specific T cell factors. The antigen also stimulates antigen-specific B cells by cross-linking their receptors, and they proliferate and express a receptor for the differentiation factor produced by the activated A cells. When the antigen-specific B cells receive the second signal differentiation factor they switch to being antibody-secreting plasma cells.

**Legend to Fig 2:**
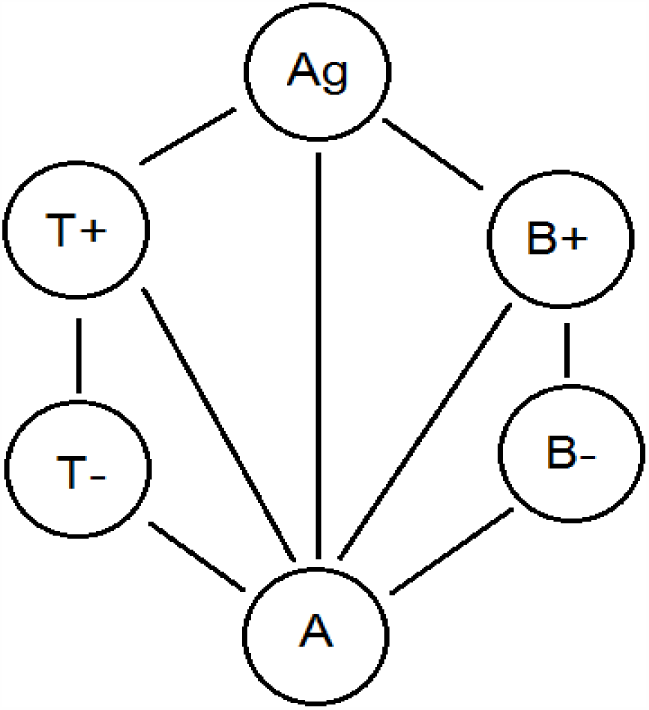
A simple idiotypic network model that includes antigen (Ag), antigen-specific T cells and B cells (T+ and B+), antiidiotypic T cells and B cells (T− and B−) and non-specific accessory cells (A cells).

## The postulated role of specific T cell factors in the induction of tolerance

Specific tolerance induction results from (a) a non-immunogenic form of an antigen stimulating T+ cells, (b) specific T cell factors from T+ cells binding to the surface of the A cells and stimulating T− cells and (c) the T− cells secreting T− specific factors and stimulating the proliferation of T+ cells. There is thus a positive feedback loop involving T+ and T− cells, taking the system to a suppressed stable steady state with elevated levels of the T+ and T− cell populations. There is co-selection (mutual selection) of the idiotypic T+ cells and the antiidiotypic T− cells. The specific T cell factors have a molecular weight of about 50,000 Daltons, and are therefore believed to be monovalent, in contrast to an IgG antibody that, with a molecular weight of 150,000, is divalent. The elevated levels of T+ and T− populations with their specific monovalent factors inhibit the stimulation of B+ and B− cells by the cross-linking of the B cell receptors.

## Conclusion

The question of whether specific T cell factors exist or not needs to be resolved. The symmetrical immune network theory (Hoffmann, 2008; Hoffmann 2010), which accounts for many aspects of the adaptive immune system, is based on the existence of such factors. In this paper, we confirm that specific T cell factors made by thymocytes do indeed exist. We submit that the fact of a confirmed experimental finding supported by a detailed theory (and vice versa) means immunology textbooks should include a section on the symmetrical immune network theory.

We propose the word “tab” as a simpler term to replace the phrase specific T cell factor.

### Materials and Methods

#### Mice

8-week old Female BALB/c mice were purchased from Jackson Labs. All mice were housed 5/cage and allowed food and water ad libitum. Animals were used for the study beginning at 10 weeks of age.

#### Immunization of mice and preparation of lymphoid extracts

5 BALB/c mice/group were immunized ×2 with 100μg/mouse KLH or BGG (iv) at 14d intervals. Mice were sacrificed 14d after the last immunization, spleen+ thymus cells pooled within groups, and cells resuspended at 3×10^8^/ml. Suspensions were sonicated at 4°C for 4min, and subjected to ultracentrifugation for 60 min at 4°C.

#### Attenuation of immune responses by sonicates from carrier immunized mice

0.3ml sonicate/mouse was infused into groups of 8 BALB/c. Control mice (8) receive no extract. 4 mice of each group of 8 mice were immunized with either DNP-KLH of DNP-BGG (100μg/mouse) in Complete Freund’s Adjuvant.

Mice were sacrificed 12d later and serum collected from all individuals. Sera were assayed in ELISA plates pre-coated with DNP-coupled albumin (100ng/well) and with HRP-anti-mouse Ig as developing Ig. Test sera were tested at 1:5, 1:20 and 1:100 dilution; only data for 1:20 dilutions are shown.

#### Statistics

Differences between groups were compared by ANOVA.

